# Endogenous oligomer formation underlies DVL2 condensates and promotes Wnt/β-catenin signaling

**DOI:** 10.1101/2024.03.07.583872

**Authors:** Senem Ntourmas, Martin Sachs, Petra Paclíková, Martina Brückner, Vítĕzslav Bryja, Jürgen Behrens, Dominic B. Bernkopf

## Abstract

Activation of the Wnt/β-catenin pathway crucially depends on polymerization of dishevelled 2 (DVL2) into biomolecular condensates. However, given the low affinity of known DVL2 self-interaction sites and its low cellular concentration it is unclear how polymers can form. Here, we detect oligomeric DVL2 complexes at endogenous protein levels, using a biochemical ultracentrifugation assay. We identify a low-complexity region (LCR4) in the C-terminus whose deletion and fusion decreased and increased the complexes, respectively. Notably, LCR4-induced complexes correlated with the formation of microscopically visible multimeric condensates. Adjacent to LCR4, we mapped a conserved domain (CD2) promoting condensates only. Molecularly, LCR4 and CD2 mediated DVL2 self-interaction via aggregating residues and phenylalanine stickers, respectively. Point mutations inactivating these interaction sites impaired Wnt pathway activation by DVL2. Our study discovers DVL2 complexes with functional importance for Wnt/β-catenin signaling. Moreover, we provide evidence that DVL2 condensates form in two steps by pre-oligomerization via high-affinity interaction sites, such as LCR4, and subsequent condensation via low-affinity interaction sites, such as CD2.

## INTRODUCTION

The Wnt/β-catenin signaling pathway promotes proliferation of stem cells, orchestrating self-renewal and regeneration of epithelial tissues in adults (Clevers et al., 2014). Deregulation of the pathway has been causally associated with severe pathologies, most prominently colorectal cancer (Clevers and Nusse, 2012). In the absence of Wnt ligands, β-catenin-dependent Wnt signaling is continuously silenced via glycogen synthase kinas 3 beta (GSK3B)-mediated phosphorylation targeting β-catenin for proteasomal degradation (Stamos and Weis, 2013). Phosphorylation of β-catenin is induced by the scaffold protein AXIN1, which interacts with β-catenin and GSK3B (Stamos and Weis, 2013). Upon binding of Wnt ligands to frizzeld receptors and low-density lipoprotein receptor-related protein 5 or 6 (LRP5/6) co-receptors, the positive Wnt pathway regulator dishevelled 2 (DVL2) interacts with frizzled and clusters beneath the plasma membrane (MacDonald and He, 2012). DVL2 clusters recruit AXIN1 and GSK3B from the cytosol, and together these proteins assemble sphere-like signalosomes at the membrane (Bilic et al., 2007). Within these signalosomes, GSK3B activity is redirected from β-catenin to LRP5/6 and finally inhibited (Bilic et al., 2007; Taelman et al., 2010). In consequence, β-catenin accumulates and can translocate into the nucleus, where it promotes transcription of its target genes (Behrens et al., 1996).

Activation of Wnt/β-catenin signaling by DVL2 crucially depends on DVL2 polymerization via its N-terminal DIX domain (Kishida et al., 1999; Schwarz-Romond et al., 2007a). However, the low auto-affinity of the DIX domain in the mid-micromolar range and the low cellular concentration of DVL2 strongly disfavor polymerization, and only the pre-clustering of DVL2 at Wnt-receptor-complexes is suggested to overcome this problem (Bienz, 2014). Mechanistically, DVL2 polymerization may support DVL2 clustering, AXIN1 recruitment and/or signalosome formation (Bilic et al., 2007; Schwarz-Romond et al., 2007a; Schwarz-Romond et al., 2007b). When DVL2 is overexpressed, DIX-mediated polymerization gives rise to microscopically visible DVL2 assemblies, which are sphere-like, membrane-free and highly dynamic (Schwarz-Romond et al., 2005). Recently, these DVL2 assemblies were characterized as phase-separated biomolecular condensates (Kang et al., 2022), which represent a functionally diverse class of membrane-free cell organelles with common biophysical properties (Banani et al., 2017; Shin and Brangwynne, 2017). In addition to the DIX domain, other parts of the protein contribute to DVL2 condensation and activity, such as the DEP domain or an intrinsically disordered region (Gammons et al., 2016; Kang et al., 2022; Vamadevan et al., 2022), suggesting that major regions of DVL2 are evolutionary optimized for condensation. Moreover, condensation of DVL2 appears to be a regulated process controlled through posttranslational modification, such as ubiquitination, and depending on its conformation, open versus close form (Lee et al., 2015; Vamadevan et al., 2022). However, there is still a controversial debate on whether DVL2 forms condensates at endogenous expression levels, as recent studies report only small DVL2 assemblies with less than 10 molecules (Kan et al., 2020), or only one big DVL2 condensate at the centrosome (Schubert et al., 2022), or about 100 condensates per cell with sizes of 0.2 to 0.5 µm (Kang et al., 2022).

In vertebrates, three DVL paralogs exist, DVL1, DVL2 and DVL3. Although overexpression of each paralog activates Wnt/β-catenin signaling, loss-of-function studies consistently report that DVL1, DVL2 and DVL3 exhibit different capabilities to transduce Wnt signals and that overexpression of one paralog does not compensate for loss of another (Lee et al., 2008; Paclikova et al., 2021). Thus, these studies point to non-redundant molecular functions of the DVL paralogs, yet, their molecular differences remain poorly understood.

Here, we report biochemical evidence for endogenous DVL2 complexes consisting of at least eight molecules, supporting the idea of DVL2 polymerization at endogenous expression levels. Using DVL2 deletion and point mutants, we mapped and characterized a low complexity region in the DVL2 C-terminus that promoted complex formation through mediating intermolecular DVL2 self-interaction. Our data suggest that these complexes most likely represent underlying substructures of DVL2 biomolecular condensates, which precede and initiate condensation. Moreover, the discovered oligomeric DVL2 complexes were of functional importance because point mutations that impaired complex formation attenuated Wnt pathway activation by DVL2.

## RESULTS

### Endogenous DVL2 forms oligomeric complexes

Performing ultracentrifugation assays, endogenous DVL2 (79 kDa) penetrated far deeper into a sucrose density gradient than AXIN1 (96 kDa) in spite of its lower molecular weight, indicating that DVL2 forms protein complexes (Fig. 1A, E). Noteworthy, most DVL2 molecules appeared to be engaged in these complexes (Fig. 1A). The complexes occurred in different cell lines (Fig. S1A), and were detectable with two, siRNA-validated antibodies (Fig. S1B, C). DVL2 (79 kDa) showed a fractionation pattern similar to thyroglobulin (669 kDa), a commercial molecular weight marker, suggesting DVL2 complex sizes of about eight molecules assuming homotypic complexes (Fig. 1A, B). As control, AXIN1 (96 kDa) showed a fractionation pattern similar to albumin (66 kDa) suggesting monomeric precipitation (Fig. 1A, B). Interestingly, the DVL2 complexes appeared to be paralog-specific because complexes were almost absent for DVL1 and DVL3 (Fig. 1C, D). The persistence of the complexes at low protein concentrations in cellular extracts indicated that they form via interaction sites with rather high affinity. Although the DIX domain is the best-characterized polymerization domain in DVL2 (Bienz, 2014), its low auto-affinity suggested that it is not involved in the formation of these complexes (Schwarz-Romond et al., 2007a). The striking difference between DVL2 and AXIN1 pointed in the same direction (Fig. 1A), since both proteins contain a functional DIX domain (Kishida et al., 1999). Consistently, a DIX domain-inhibiting point mutation (DVL2 M2) (Schwarz-Romond et al., 2007a) did not affect DVL2 complexes (Fig. 1C, D).

**Fig. 1.**
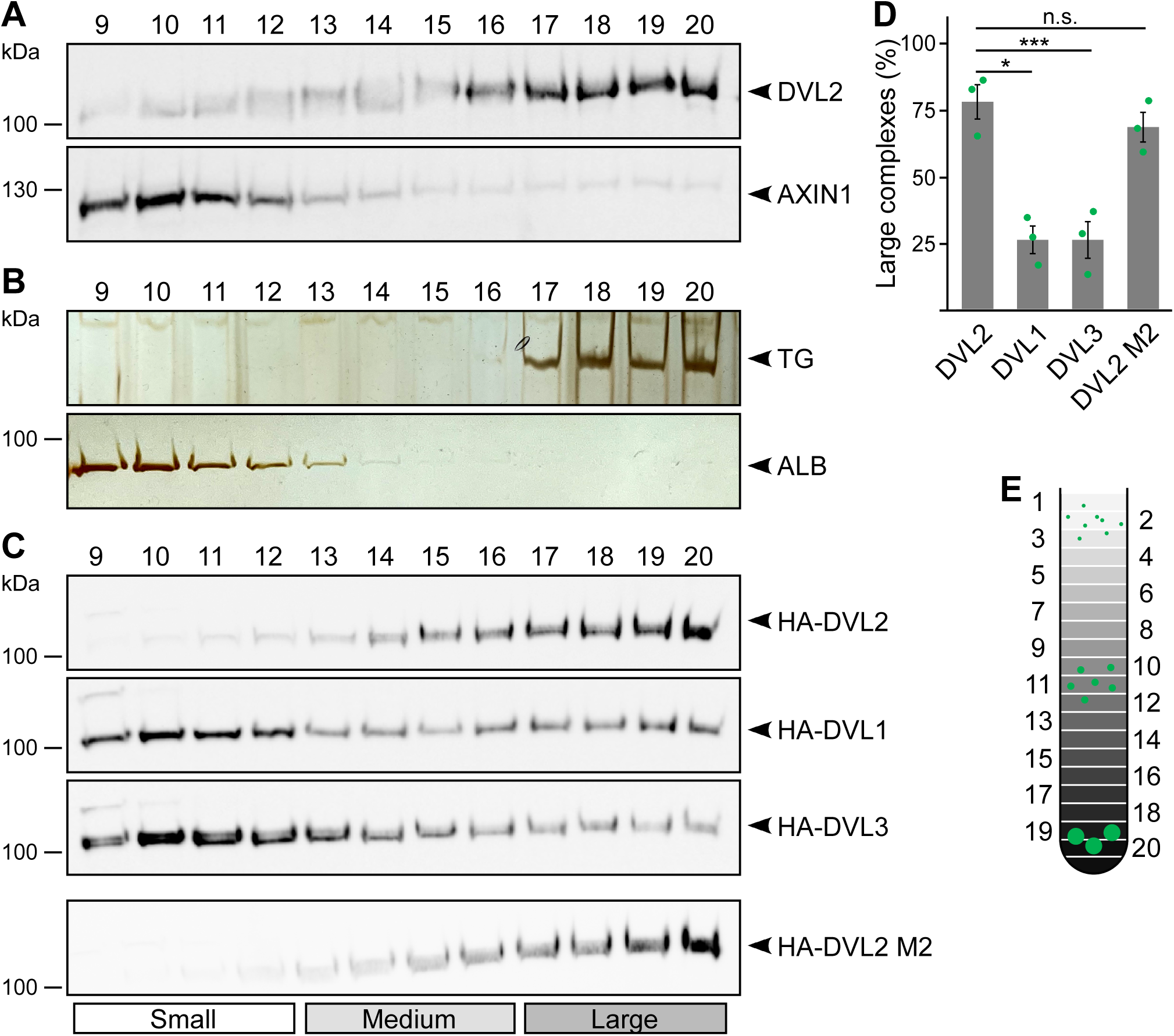
Endogenous DVL2 forms paralog-specific oligomeric complexes. **A**, **C** Western blotting for indicated endogenous (**A**) or transiently expressed proteins, detected with anit-HA antibodies (**C**) after fractionation of HEK293T cell lysates via sucrose density ultracentrifugation. **B** Silver staining of thyroglobulin (TG) or albumin (ALB) after sucrose density ultracentrifugation of purified proteins. **A**-**C** shows one out of at least three representative experiments. Analyzed fractions are indicated above the blots according to **E**. **D** Amount of the protein that was engaged in large complexes (see label Large in **C**), relative to the cumulative protein amount detected in all investigated fractions, as determined by 2D densitometry analysis of protein bands from three independent experiments as in **C** (n=3). Results are mean +-SEM, * p<0.05, *** p<0.001 (Student’s *t*-test) **E** Schematic representation of the ultracentrifugation assay, illustrating distribution of proteins of different sizes (green) within numbered fractions of a sucrose gradient form low (light grey) to high density (black).

### A low complexity region in the C-terminus promotes DVL2 complexes

Deletion of the DEP domain (construct ΔDEP) decreased DVL2 complexes, in line with its reported function in DVL2-DVL2 interaction (Gammons et al., 2016). However, additional deletion of the remaining C-terminus (construct 1-418) markedly reduced them further, demonstrating a strong contribution of the deleted residues 521-736 to complex formation (Fig. 2A-C). Importantly, decreased protein complexation of DVL2 ΔDEP and 1-418 in ultracentrifugation experiments conspicuously correlated with decreased formation of condensates in immunofluorescence-based assays (Fig. 2D, E) and decreased Wnt pathway activation in reporter assays (Fig. 2F). Therefore, we hypothesized that the DVL2 complexes may be important for signaling activity. To identify candidate regions for a more precise mapping within residues 521-736, we used the SEG algorithm predicting low-complexity regions (Wootton and Federhen, 1993), the TANGO algorithm predicting aggregation (Fernandez-Escamilla et al., 2004) and protein alignments (Sievers et al., 2011). We identified four low-complexity regions (LCR1-4), which are associated with protein assembly (Martin et al., 2020; Martin and Mittag, 2018), one potential aggregation site embedded in LCR4, and two domains whose evolutionary conservation may point to functional importance (CD1-2, Fig. S1D). Since deletion of residues 521-736 showed strong effects when combined with the DEP deletion (1-418 vs ΔDEP, Fig. 2), we performed the following mapping in the ΔDEP context. Given the good correlation between complexes and condensates (Fig. 2C, E), we decided to use immunofluorescence-based analysis of condensates for mapping, as it is more convenient than density gradient ultracentrifugation. Upon individual deletion of the six identified regions LCR1-4 and CD1-2, only deletion of LCR4 and CD2 decreased condensate formation and Wnt pathway activation of DVL2 ΔDEP (Figs. 3A and S2A-C). Combined deletion of LCR4 and CD2 increased the effect (Figs. 3A and S2A-C). We, therefore, consider the two adjacent regions as one functional unit, hereafter referred to as condensate forming region (CFR). DVL2 ΔDEP-ΔCFR exhibited a marked decrease in the number of cells with condensates (Fig. 3A-C), in the number of condensates per cell (Figs. 3D and S2D-G) and in Wnt pathway activation (Fig. 3E), similar to 1-418. Importantly, individual or combined fusion of LCR4 and CD2 to 1-418 sufficed to induce condensation (Figs. 3A-D, S2D-G and S3A, B) and Wnt pathway activation (Figs. 3E and S3C, D), rendering 1-418+CFR as active as ΔDEP. Mutational inactivation of the DIX domain (1-418+CFR M2) abolished condensates demonstrating that the DIX domain is required for CFR-mediated condensates (Fig. S3A, B). Interestingly, the investigated DVL2 mutant proteins predominantly formed nuclear condensates in contrast to the cytosolic condensates of WT DVL2, most likely, because a nuclear export signal (Fig. 2A) was deleted in these mutants (Fig. 3A). However, investigating only cells with cytosolic condensates (Fig. S3E, F) revealed similar differences between the DVL2 mutants as were observed when investigating mainly cells with nuclear condensates (Fig. 3C and S3B), suggesting that the detected differences are not due to nuclear localization but reflect the overall condensation capacity of the DVL2 mutants. Moreover, fusion of CFR to the isolated DVL2 DIX or AXIN1 DAX domain sufficed to trigger condensation, which was prevented by M2/M3 mutation of the DIX/DAX domain (Fig. S3D, G, H). Thus, loss- and gain-of-function experiments identified CFR as the crucial region for condensation and Wnt pathway activation within the DVL2 C-Terminus, which functionally cooperates with the DIX domain.

**Fig. 2.**
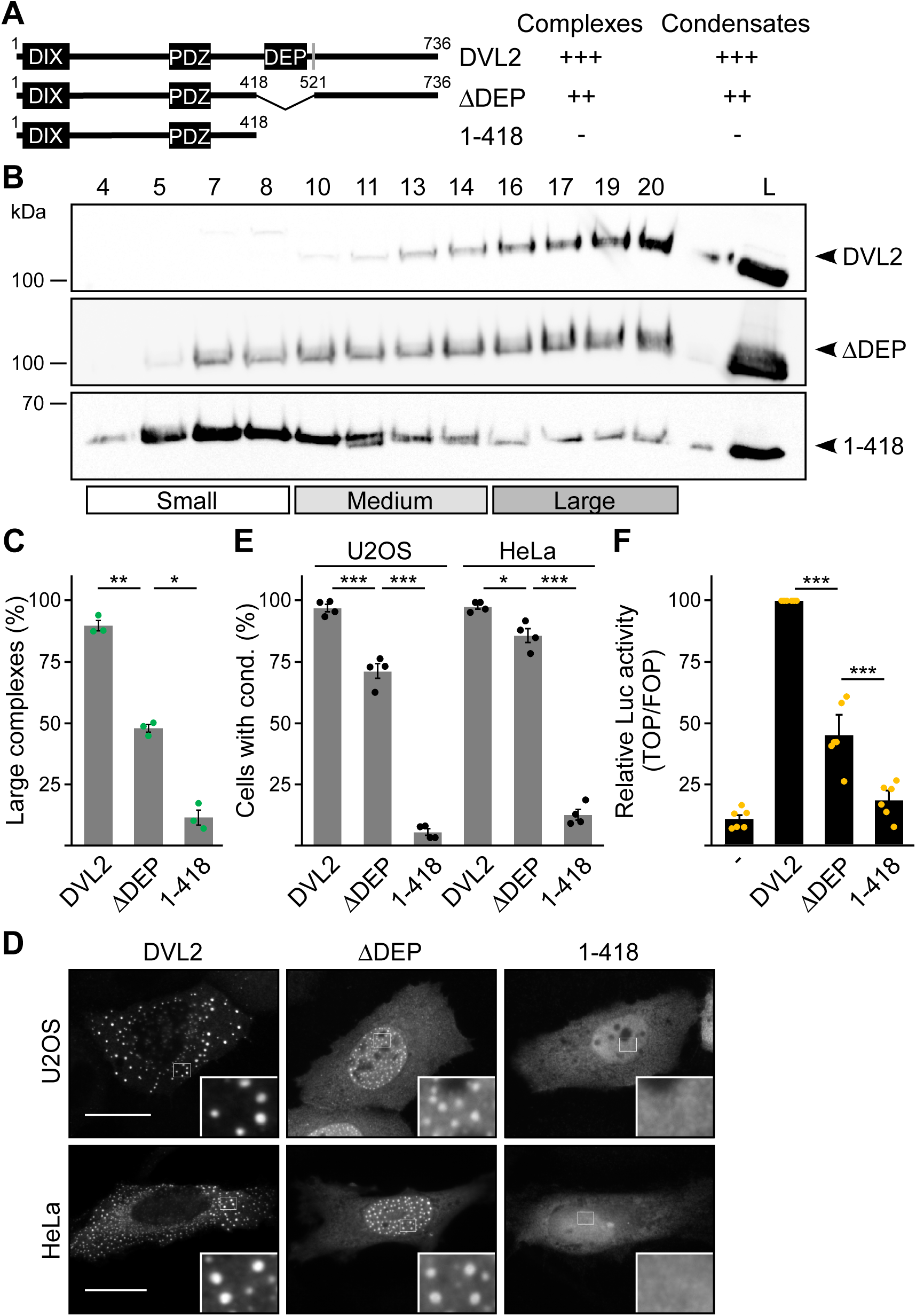
The DVL2 C-terminus promotes complexes, condensates and activity. **A** To scale schemes of DVL2 constructs with the DIX, the PDZ and the DEP domain. A nuclear export signal is highlighted in grey (Itoh et al., 2005). Indicated complexation and condensation summarizes the findings in **B**-**E**. **B** Western blotting for indicated transiently expressed proteins bevor (L) and after fractionation (4-20) of HEK293T cell lysates via sucrose density ultracentrifugation. **C** Percentage of the protein that was engaged in large complexes as specified in **B** (n=3, refer to the legend to Fig. 1D for more details). **D** Immunofluorescence of indicated HA-tagged proteins in transiently transfected U2OS and HeLa cells. Scale bars: 20 µm. Insets are magnifications of the boxed areas. Interestingly, DEP domain deleted constructs frequently showed nuclear condensates in contrast to the cytosolic condensates of full length DVL2, which is most likely explained by the deletion of a nearby nuclear export signal (see **A**) and which still allowed determining differences in condensation capacity. **E** Percentage of cells with condensates out of 1200 transfected cells from four independent experiments as in **D** (n=4). **F** Relative luciferase activity reporting β-catenin-dependent transcription in HEK293T cells expressing the indicated constructs (n=6). **C**, **E**, **F** Results are mean +-SEM, * p<0.05, ** p<0.01, *** p<0.001 (Student’s *t*-test).

**Fig. 3.**
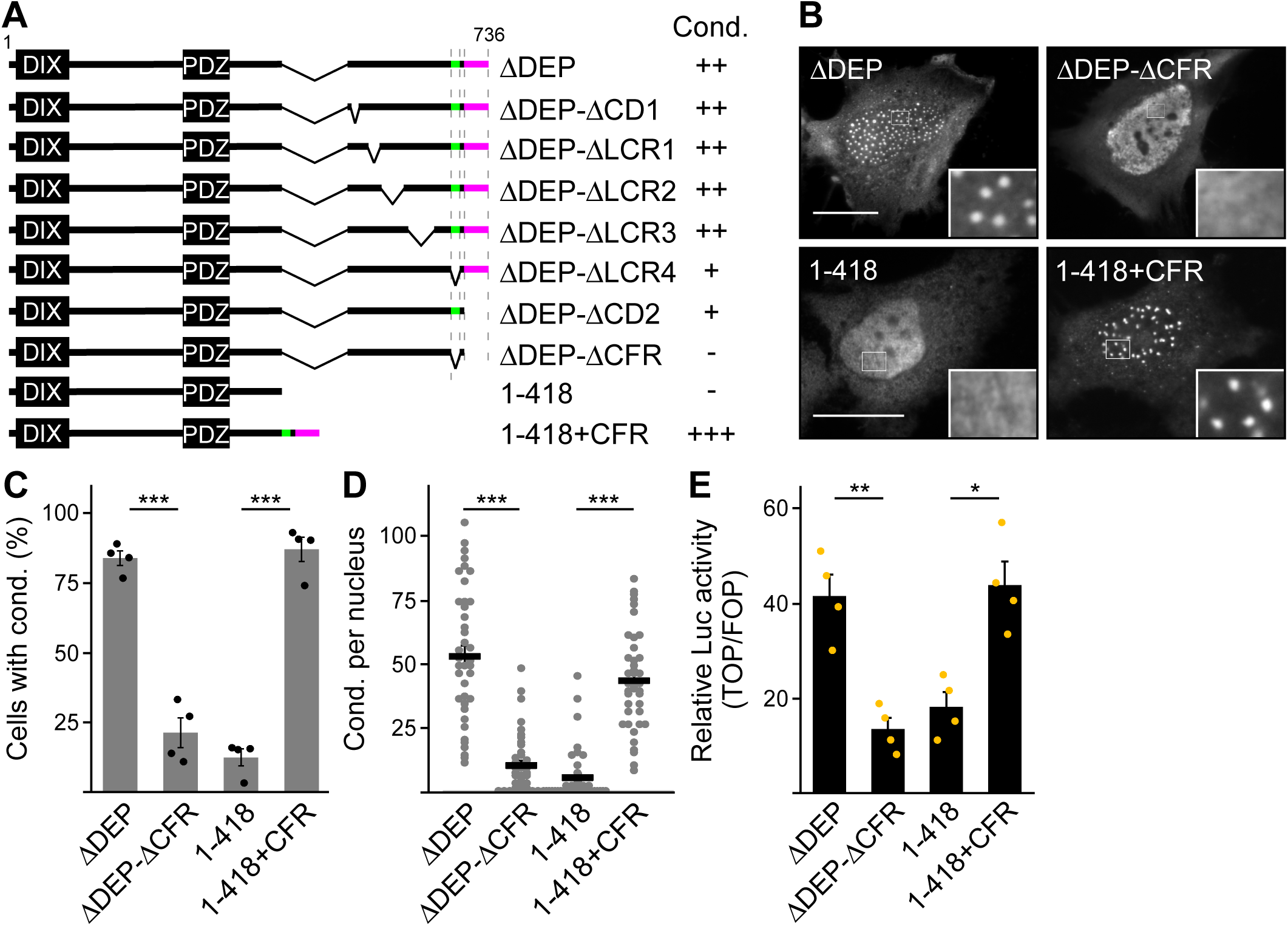
A 58 aa C-terminal region promotes DVL2 condensates and activity. **A** To scale schemes of DVL2 constructs. Indicated condensation (Cond.) summarizes the findings in Fig. S2B, and the identified crucial regions are highlighted in green (LCR4) and magenta (CD2). **B** Immunofluorescence of indicated HA-tagged proteins in transiently transfected HeLa cells. Scale bars: 20 µm. Insets are magnifications of the boxed areas. **C** Percentage of cells with condensates out of 1200 transfected cells from four independent experiments as in **B** (n=4). **D** Automated quantification of condensate number per nucleus by the Icy Spot Detector (Olivo-Marin, 2002) in 40 cells from four independent experiments as in **B** (n=40). **E** Relative luciferase activity reporting β-catenin-dependent transcription in HEK293T cells expressing the indicated constructs (n=4). **C**-**E**, Results are mean +-SEM, * p<0.05, ** p<0.01, *** p<0.001 (Student’s *t*-test).

Next, we investigated whether CFR contributes to the formation of DVL2 complexes detected by density gradient ultracentrifugation. Indeed, CFR deletion from ΔDEP (ΔDEP-ΔCFR) and fusion with 1-418 (1-418+CFR) decreased and increased complexes, respectively (Fig. 4A-D). Moreover, replacing the respective sequence in DVL1 with the CFR of DVL2 (DVL1-CFR^DVL2^) promoted complex formation (Fig. 4E, F). Surprisingly, only LCR4 but not CD2 was required for complex formation when deleted from ΔDEP or mediated complex formation when fused to 1-418 (Fig. 4A-D), although both parts were required for condensate formation (Fig. S2B). Of note, LCR4 is not well conserved in DVL1 and DVL3, which do not form complexes (Figs. S1D and 1C). Together, our data revealed a bipartite, 58 amino acid region at the very C-Terminus of DVL2 consisting of a low complexity region (LCR4) that promotes complexes and condensates and a conserved domain (CD2) that only promotes condensates and is dispensable for complexes.

**Fig. 4.**
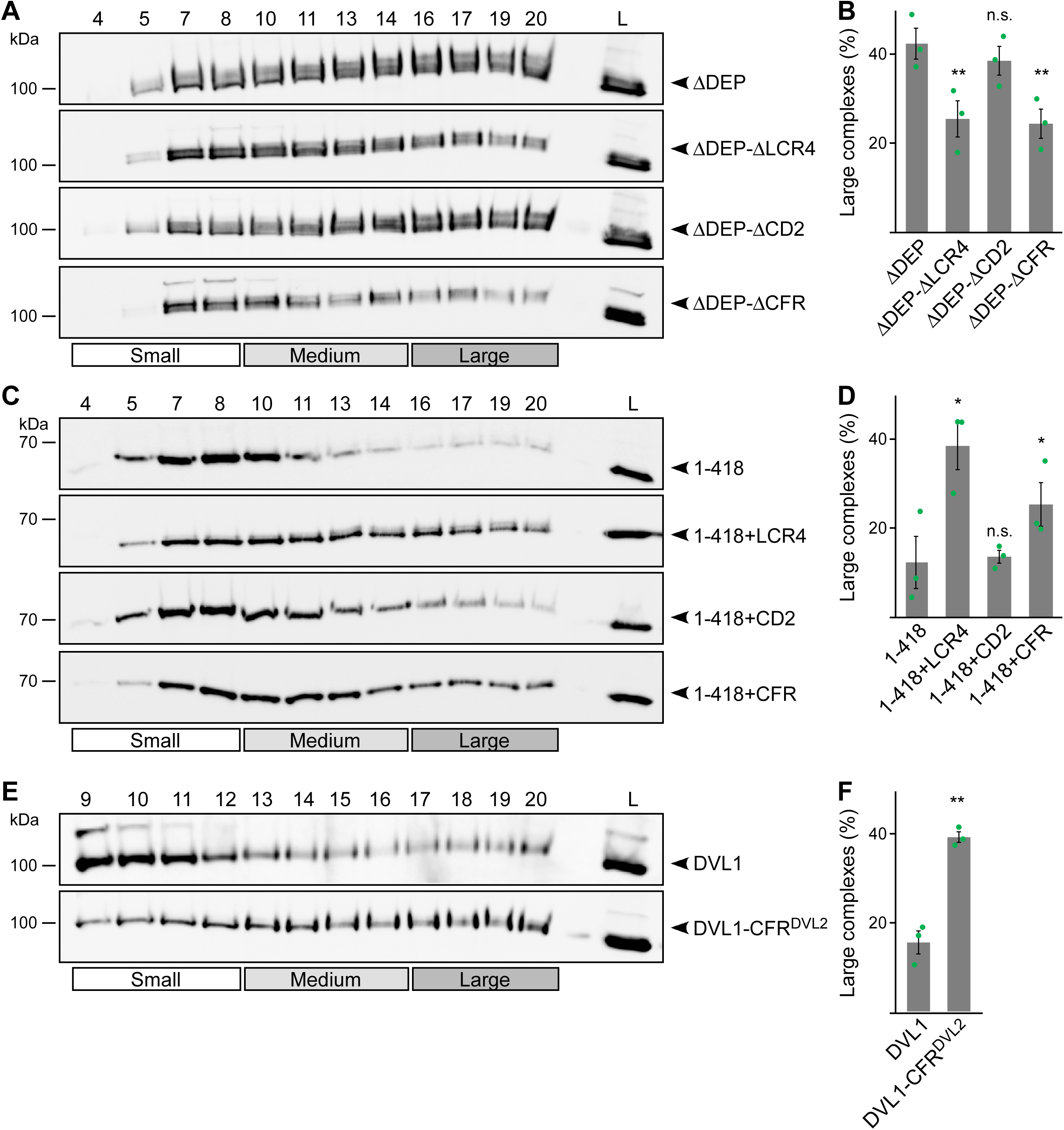
LCR4 mediates complex formation. **A**, **C**, **E** Western blotting for indicated transiently expressed proteins bevor (L) and after fractionation (4-20) of HEK293T cell lysates via sucrose density ultracentrifugation. **B**, **D**, **F** Percentage of the protein that was engaged in large complexes as specified in **A**, **C** or **E** (n=3, refer to the legend to Fig. 1D for more details). Results are mean +-SEM, * p<0.05, ** p<0.01 (Student’s *t*-test).

### LCR4 and CD2 cooperatively mediate DVL2 self-interaction

Given the role of the CFR parts LCR4 and CD2 in complex and condensate formation of DVL2, we speculated that they might directly mediate DVL2-DVL2 interaction. To analyze the role of a putative aggregation site (aggregon) that was predicted by the TANGO algorithm within LCR4 (Fig. S1D), we designed mutations of two key valine residues (VV-AA) predicted to prevent aggregation (Figs. 5A and S5A). LCR4 VV-AA mutation of DVL2 ΔDEP decreased condensate formation as efficiently as LCR4 deletion (Fig. 5B, C), and fusion of VV-AA-mutated LCR4 failed to increase condensate formation of 1-418, in contrast to WT LCR4 (Fig. S5B, C). In CD2, the TANGO algorithm did not predict aggregation sites. However, we detected interspersed phenylalanine residues in CD2 (Fig. 5A), which might promote interaction through stacking of their aromatic rings, acting as “stickers” of unstructured regions, as recently described (Martin et al., 2020). CD2 FF-AA mutation decreased condensate formation and Wnt pathway activation as efficiently as CD2 deletion (Figs. 5B, C and S5D), and CD2 FF-AA fusion failed to increase condensate formation or Wnt pathway activation, in contrast to WT CD2 (Fig. S5E-G). Our data suggest that CFR is a bipartite protein interaction site for self-association of DVL2. In line with this, the isolated CFR, which exhibited a homogeneous cellular distribution when expressed alone, accumulated within DVL2 condensates upon co-expression of both proteins (Fig. 5D, E), suggesting CFR-DVL2 interaction. CFR VV-AA FF-AA mutation markedly reduced this CFR-DVL2 co-localization (Fig. 5D, E). Moreover, DVL2 CFR did not co-localize with DVL1 or DVL3 condensates (Fig. S5H), consistent with the low conservation of the LCR4 aggregon in these paralogs (Fig. S1D). Notably, co-expression of the isolated CFR inhibited Wnt pathway activation by DVL2 in a dosage-dependent manner, which was attenuated by CFR FF-AA mutation (Fig. 5 F). In this experiment, free CFR might weaken DVL2-DVL2 interaction by saturating the interaction surface of the CFR within DVL2. Together, our data provide evidence that CFR mediates intermolecular DVL2 self-interaction via an aggregation site in LCR4 and phenylalanine stickers in CD2. Since individual deletion of LCR4 or CD2 was sufficient to impair condensates (Fig. S2B), both interaction sites most likely cooperate to drive condensate formation of DVL2.

**Fig. 5.**
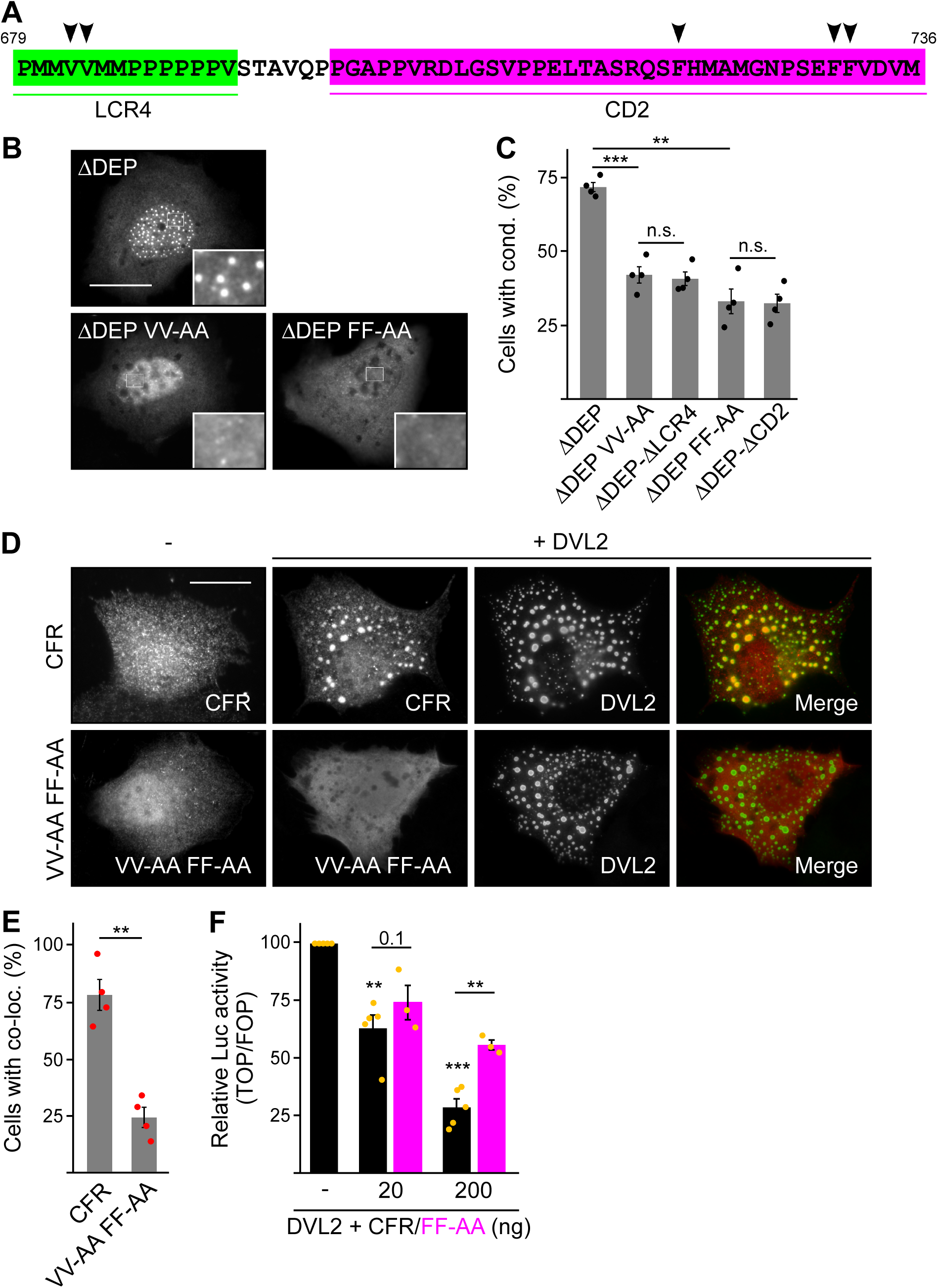
LCR4 and CD2 cooperatively mediate DVL2-DVL2 self-interaction. **A** CFR amino acid sequence with highlighted LCR4 (green) and CD2 (magenta). Arrowheads point to residues potentially mediating protein-protein interaction. **B** Immunofluorescence of indicated HA-tagged proteins in transiently transfected U2OS cells. Scale bars: 20 µm. Insets are magnifications of the boxed areas. **C** Percentage of cells with condensates out of 1200 transfected cells from four independent experiments as in **B** (n=4). **D** Immunofluorescence of indicated proteins in U2OS cells, which were transfected with Flag-CFR or the Flag-CFR VV-AA FF-AA mutant either alone or in combination with DVL2. Scale bar: 20 µm. **E** Percentage of cells exhibiting co-localization of CFR or CFR VV-AA FF-AA with DVL2 out of 1200 transfected cells from four independent experiments as in **D** (n=4). **F** Relative luciferase activity reporting β-catenin-dependent transcription in HEK293T cells transfected with DVL2 alone or together with rising amounts of either CFR or the CFR FF-AA mutant (black bars [CFR] n=5, magenta bars [CFR FF-AA] n=3). **C**, **E**, **F** Results are mean +-SEM, ** p<0.01, *** p<0.001 (Student’s *t*-test).

### DVL2 CFR promotes phase separation

While 1-418 and DVL2 DIX exhibited a homogenous cellular distribution, fusion of CFR induced spherical, microscopically visible condensates of 1-418+CFR and DIX+CFR (Figs. 3B and S3G). To study the nature of CFR-mediated condensates, we treated cells expressing DVL2, 1-418+CFR or DIX+CFR with a hypoosmolar buffer (osmotic shock) or with the bivalent alcohol 1,6-hexandiol, as previously done in biomolecular condensate research to challenge phase separation (Nott et al., 2015; Ribbeck and Gorlich, 2002). Both treatments significantly decreased condensates of all three studied proteins within 3 min for osmotic shock and 1 h for 1,6-hexandiol (Fig. 6A-C, and movies M1 and M2). These findings are consistent with the transition from a two-phase state of condensates with high protein concentration and surrounding spaces with low protein concentration to a one-phase state of homogenous protein distribution (Fig. 6A). We concluded that CFR indeed induced phase separation to promote 1-418+CFR and DIX+CFR condensates, in line with the fact that WT DVL2 was shown to undergo phase separation (Kang et al., 2022),. Moreover, fusion of an AXIN1-derived, sequencewise-nonrelated condensate-forming region (CFR^AX^, see Fig. S4 for details) to DVL2 ΔDEP-ΔCFR (ΔDEP-ΔCFR+CFR^AX^) restored condensate formation to the level of ΔDEP (Fig. 6D-F), indicating that CFR^AX^ compensates for loss of CFR^DVL2^. More importantly, CFR^AX^ fusion (ΔDEP-ΔCFR+CFR^AX^) also rescued the decreased Wnt pathway activation of ΔDEP-ΔCFR compared to ΔDEP (Fig. 6G), suggesting that it is indeed the CFR phase-separating activity that is crucial for signaling.

**Fig. 6.**
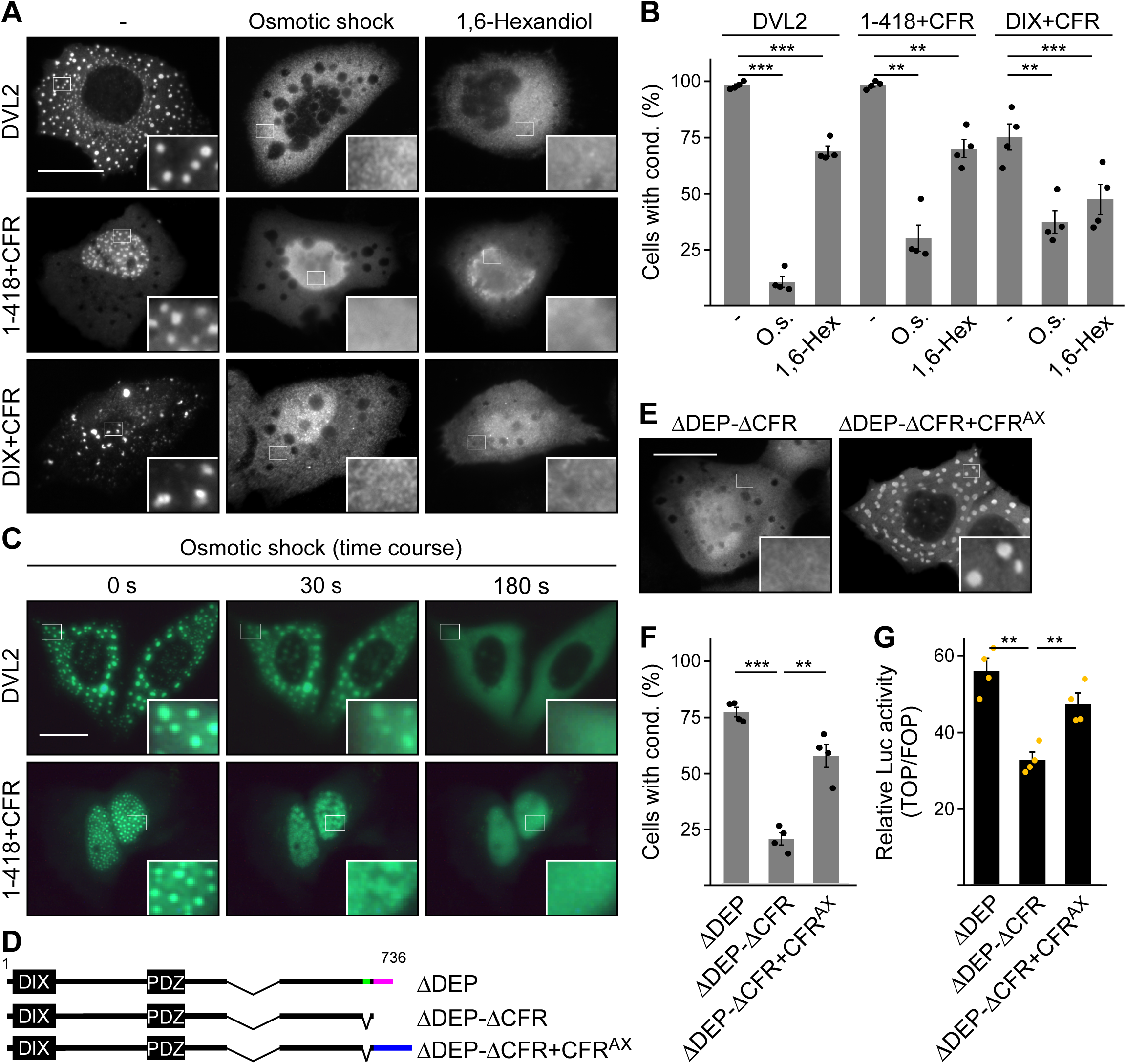
CFR-induced condensates form via phase separation. **A**, **E** Immunofluorescence of indicated HA-tagged proteins in transiently transfected U2OS cells, which were untreated, exposed to osmotic shock for 3 min, or treated with 1 µM 1,6-hexandiol for 1 h. Scale bars: 20 µm. Insets are magnifications of the boxed areas. **B**, **F** Percentage of cells with condensates out of 1200 transfected cells from four independent experiments as in **A** or **E** (n=4). **C** Fluorescence of indicated GFP-tagged proteins in transiently transfected, alive U2OS cells at different time points of osmotic shock treatment. Scale bar: 20 µm. Insets are magnifications of the boxed areas. **D** To scale schemes of DVL2 constructs. LCR4, CD2 and CFR^Ax^ are highlighted in green, magenta and blue, respectively. **G** Relative luciferase activity reporting β-catenin-dependent transcription in HEK293T cells expressing the indicated constructs (n=4). **B**, **F**, **G** Results are mean +-SEM, ** p<0.01, *** p<0.001 (Student’s *t*-test).

### DVL2 CFR contributes to Wnt pathway activation

In order to determine the impact of CFR within full-size DVL2, we mutated the four crucial residues identified in CFR in DLV2 (DVL2 VV-AA FF-AA). This shifted the DVL2 complexes in ultracentrifugation assays from large to smaller sizes, similar to deletion of CFR (Fig. 7A, B), and reduced the formation of condensates by about 50% (Fig. 7C, D). Importantly, DVL2 VV-AA FF-AA exhibited more than 50% reduced Wnt pathway activation compared to WT (Fig. 7E), with similar expression of both constructs (Fig. S5I). The DVL2 variants were transiently expressed on top of endogenous DVL1/2/3 in this experiment. In addition, we used *DVL1/2/3* knockout cells, as they represent an elegant system to study Wnt pathway activation upon re-expression of DVL2 variants without any interference of endogenous WT DVL (Paclikova et al., 2017). In these cells, overexpression of DVL2 VV-AA FF-AA almost completely failed to activate the pathway and was as inactive as the DIX domain M2 mutant (Schwarz-Romond et al., 2007a), which can be considered as the gold standard for DVL2 inhibiting point mutations (Fig. 7F). In addition, we used *DVL1/2/3* knockout cells with additional knockout of *RNF43* and *ZNRF3* (DVL tKO+), which allowed higher pathway activation upon DVL2 overexpression (Fig. S5J), as the DVL2-activating receptors were no longer degraded through RNF43 and ZNRF3 (Hao et al., 2012; Paclikova et al., 2021). Also in these cells, DVL2 VV-AA FF-AA exhibited markedly impaired pathway activation as compared to WT (Fig. S5J). Finally, we re-expressed DVL2 variants at close to endogenous levels in *DVL1/2/3* knockout cells to rescue Wnt-induced pathway activation, which was disrupted through DVL knockout (Figs. 7G and S5K). While re-expression of WT DVL2 resulted in a complete rescue as compared to WT cells, DVL2 VV-AA FF-AA was significantly impaired and as inactive as DVL2 M2 in this assay (Fig. 7G). The VV-AA FF-AA mutation inhibited complexation, condensation and Wnt pathway activation as efficiently as CFR deletion (Fig. 7A-F), strongly indicating that it is specifically the interaction activity of CFR through the aggregon and the phenylalanine stickers that is required for signaling. A comparison between the VV-AA FF-AA mutation and the established M2 mutation showed on average about 65% and 80% reduction of Wnt pathway activation as compared to WT, respectively (Figs. 7E-G and S5K), suggesting that DVL2 CFR markedly contributes to Wnt pathway activation. Consistently, we observed strong significant correlations of CFR-mediated condensation and complexation with Wnt pathway activation for the DVL2 deletion constructs used in our study (Fig. 7H).

**Fig. 7.**
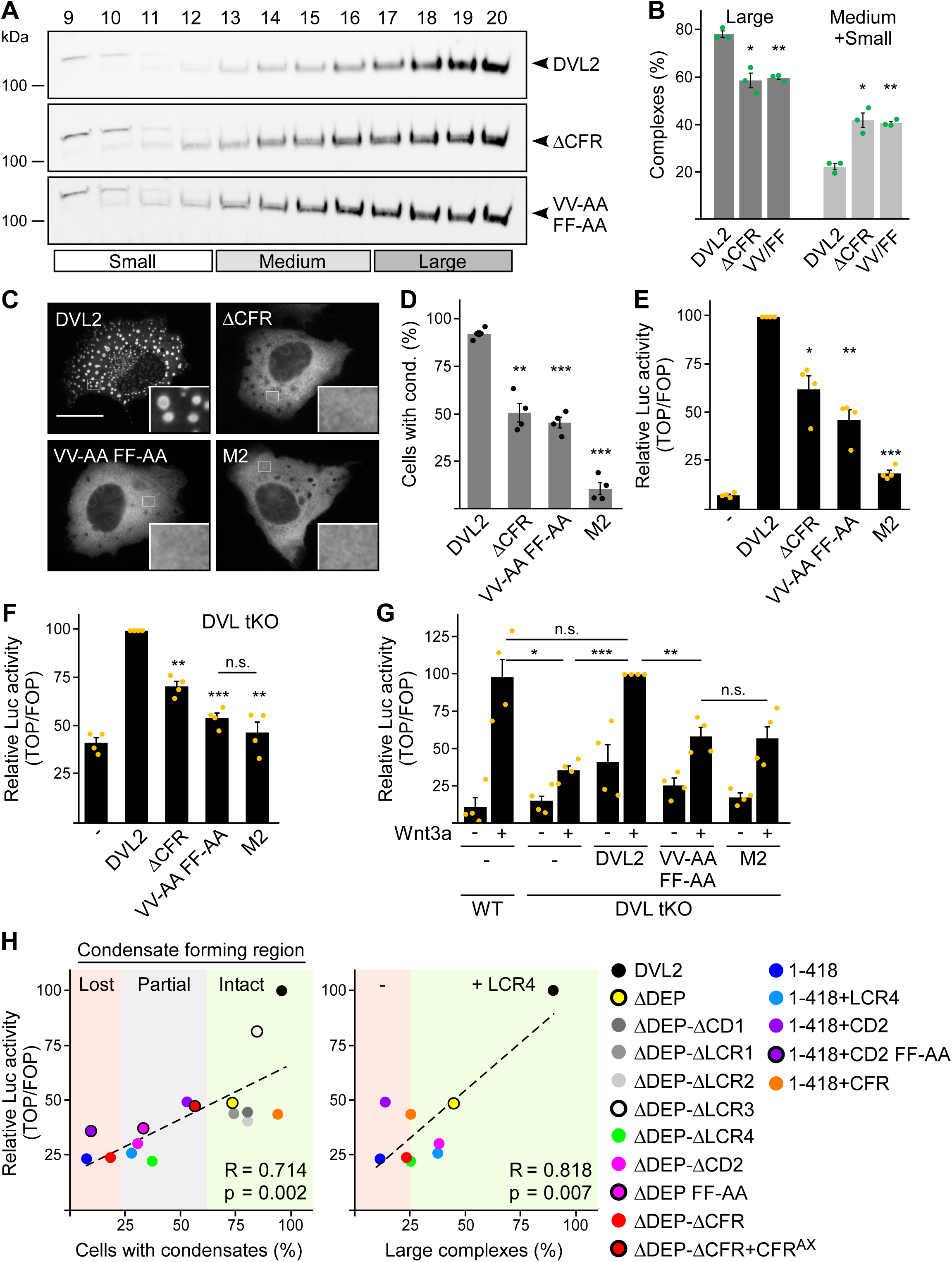
DVL2 CFR is crucial for Wnt signaling activity. **A** Western blotting for indicated proteins, which were transiently expressed in HEK293T cells, after fractionation of cell lysates via sucrose density ultracentrifugation (see Fig. 1E). **B** Percentage of the protein that was engaged in large complexes or in medium/small complexes, as specified in **A** (n=3, refer to the legend to Fig. 1D for more details). **C** Immunofluorescence of indicated HA-tagged DVL2 proteins in transiently transfected U2OS cells. Scale bar: 20 µm. Insets are magnifications of the boxed areas. **D** Percentage of cells with condensates out of 1200 transfected cells from four independent experiments as in **C** (n=4). **E**-**G** Relative luciferase activity reporting β-catenin-dependent transcription in U2OS cells (**E**), in T-REx cells with *DVL1/2/3* knockout (**F**, DVL tKO) and in T-REx WT and DVL tKO cells (**G**), which were transiently transfected and treated with Wnt3a conditioned medium, as indicated (n=4). **B**, **D**-**G** Results are mean +-SEM, * p<0.05, ** p<0.01, *** p<0.001 (Student’s *t*-test). **H** Plotting of Wnt pathway activation (y-axis) against either condensation (x-axis; left side) or complexation (x-axis; right side) for indicated DVL2 WT and mutant proteins. Correlation strength and significance are indicated by the Pearson‘s correlation coefficient R and the p value, respectively. Note that condensation correlates with whether CFR is intact (LCR4 and CD2 intact, green), partially intact (either LCR4 or CD2 intact, grey) or lost (neither LCR4 nor CD2 intact, red), and that the presence of LCR4 determines complexation. The plots summarize data that were shown before within this study.

## DISCUSSION

Here, we provided strong biochemical evidence that endogenous DVL2 forms oligomeric complexes (Fig. 1A, B), supporting the idea that DVL2 assemblies exist at physiologic protein levels. Although we investigated several scaffold proteins with various endogenous interactors in this assay, we only observe such complexes for DVL2 and AXIN2 and they were specifically associated with aggregating protein sequences (Bernkopf et al., 2019; Miete et al., 2022), indicating that most types of protein-protein interactions are not preserved in this assay. Furthermore, overexpressed DVL2 did not exhibit reduced complex sizes, as one would have expected, if limited endogenous interactors had been part of the complexes. Therefore, we think that the detected complexes most likely reflect homotypic DVL2 assemblies, which would then be about eight molecules in size (Fig. 1A, B). The size of the complexes was reminiscent of previously described endogenous DVL2 oligomers identified via TIRF imaging (Kan et al., 2020). Through deletion analysis, we discovered a 14 aa long, low complexity region (LCR4) in the DVL2 C-terminus mediating complex formation (Fig. 4A-D) and condensate formation (Fig. S2B). An adjacent 38 aa long, evolutionary conserved domain (CD2) was required for the formation of DVL2 condensates (Fig. S2B), and we conceptually combined LCR4 and CD2 as condensate forming region (CFR). Molecularly, an aggregon in LCR4 and phenylalanine residues in CD2 mediated DVL2 assembly (Figs. 7A-D and S5A-G). The latter may promote protein interaction via sticking of their aromatic rings, as previously described for phase-separating proteins (Martin et al., 2020). Co-localization of the isolated CFR with DVL2 indicated that CFR may mediate DVL2-DVL2 interaction, and this was prevented by specific point mutations targeting the aggregon in LCR4 and the phenylalanine stickers in CD2 (Fig. 5D, E). Treatments that challenge phase separation diffused CFR-induced condensates (Fig. 6A-C), in line with a recent report showing that DVL2 condensates form via phase separation (Kang et al., 2022). Importantly, point mutations that inhibit CFR self-interaction markedly attenuated Wnt pathway activation by DVL2 (Fig. 7E-G and S5K). Especially in *DVL1/2/3* triple knockout cells, the DVL2 CFR point mutant VV-AA FF-AA was as signaling deficient as the DIX domain M2 mutant (Fig. 7F, G), which is frequently used for DVL2 inactivation (Schwarz-Romond et al., 2007a). In these cells, mere DVL2 VV-AA FF-AA overexpression almost completely failed to activate the pathway (Fig. 7F) and its re-expression at close to endogenous levels only poorly rescued activation of the pathway through Wnt ligands, which was disrupted by the DVL knockout (Figs. 7G and S5K). We thus identified a novel DVL2 region that promotes complex formation, condensate formation, and Wnt pathway activation.

Our study provides deeper insights into the assembly of multimeric DVL2 condensates. The complexes detected by ultracentrifugation showed sizes of about eight DVL2 molecules, which were irrespective of expression levels as they were similar between endogenous and overexpressed DVL2 (Fig. 1A-C). Therefore, the complexes cannot be identical with the known microscopically visible condensates, containing thousands of molecules. These oligomeric complexes thus existed in parallel to the condensates or constituted a condensate substructure. Analysis of mutant DVL2 proteins revealed that DVL2 contains domains that promote both complexes and condensates, such as LCR4 and DEP (Figs. 2B-E, 4A-D and S2B), and domains that promote only condensates but not complexes, such as CD2 and DIX (Figs. 1C, 4A-D, 7C and S2B). However, all DVL2 mutations that reduced complexes also affected condensates. Therefore, we suggest that the oligomeric complexes are required for and possibly represent substructures of the multimeric condensates (Fig. 8). As the stability of complexes at low protein concentrations, e.g. in cellular extracts, indicated a rather high affinity of the underlying interaction sites (Fig. 1A), we hypothesize that complex formation via LCR4 and DEP precedes further assembly of these substructures into condensates via CD2 and DIX (Fig. 8). Our proposed two-step model of DVL2 condensate formation is in line with and supportive of the emerging stickers-model for the formation of biomolecular condensates (Choi et al., 2020). According to this model, oligomerization via one interaction site can drive subsequent condensate formation by increasing the valence of another interaction site (=sticker) of the oligomer compared to a monomer (Choi et al., 2020). The proposed two-step model would increase the options for regulating polymerization of DVL2 by modulating the oligomerization step or the condensate formation step (Fig. 8). DVL2 polymerization is crucial for Wnt signal transduction. However, the low DVL2 concentration and the low DIX-DIX affinity strongly disfavor polymerization, and clustering of DVL2 at activated Wnt-receptor-complexes is suggested to overcome this limitation (Bienz, 2014). Pre-oligomerization of DVL2 via high-affinity interactions as identified in the ultracentrifugation assay might facilitate Wnt-induced polymerization.

**Fig. 8.**
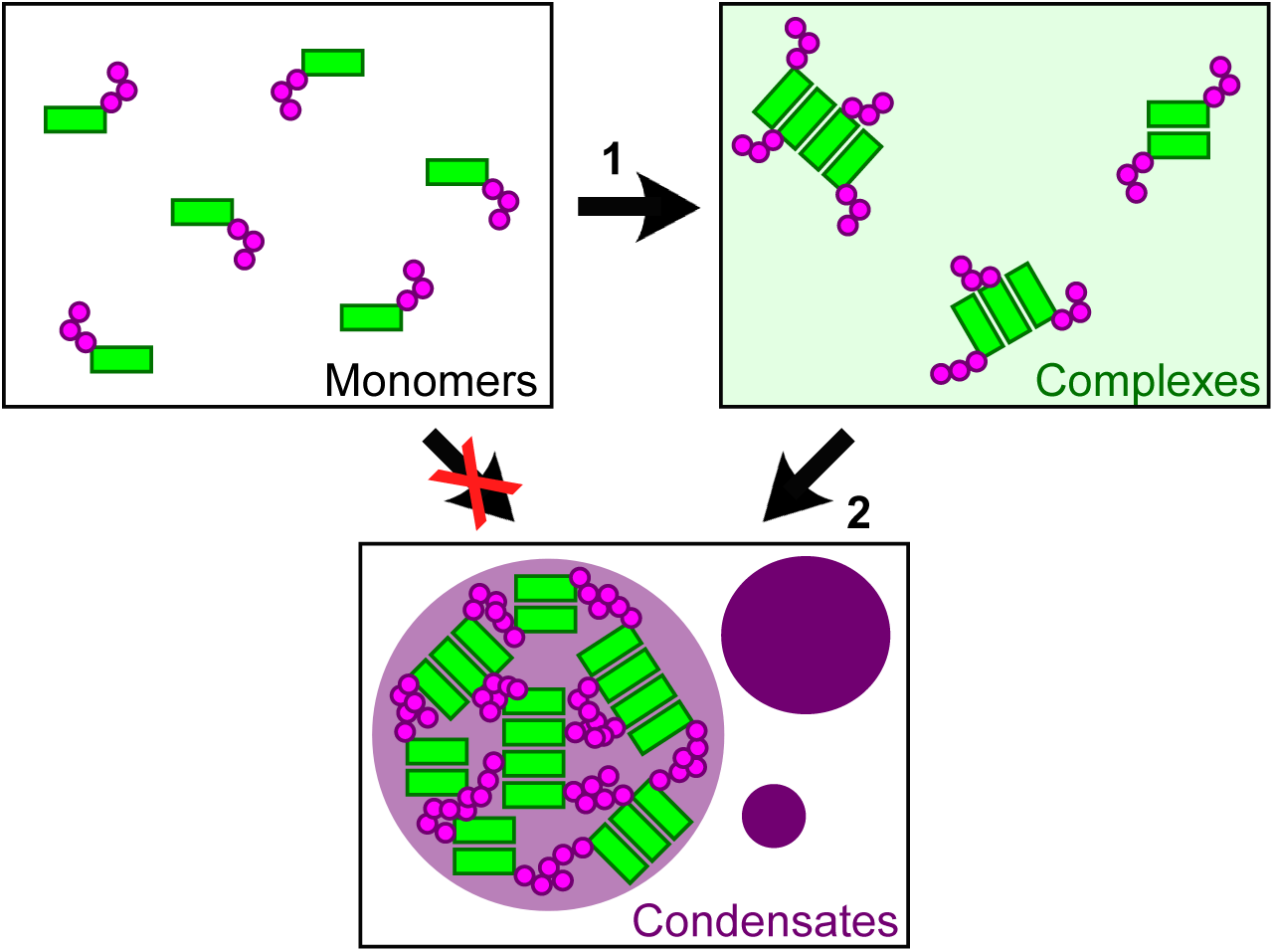
Two-step model of DVL2 condensate formation. Schematic illustration of DVL2 domains mediating the formation of oligomeric complexes, such as LCR4 and the DEP domain (green rectangles) and of sticker domains mediating condensate formation, such as CD2 and the DIX domain (magenta circles). Oligomerization into complexes (1) (light green background, right) increases the valence of stickers in the complexes as compared to the monomers, which allows to overcome the low affinity of the isolated stickers and drives subsequent formation of condensates (2) (purple, bottom) by the multivalent stickers, according to the emerging stickers-model (Choi et al., 2020).

Furthermore, the discovered C-terminal self-interaction site CFR may contribute to the regulation of DVL2 condensates through conformational changes. Binding of the very C-terminus of DVL2 to its PDZ domain results in a closed protein conformation (Lee et al., 2015). It has been suggested that this closed conformation limits the accessibility of the N-terminal intrinsically disordered region that promotes condensate formation, thereby suppressing DVL2 condensates (Kang et al., 2022; Vamadevan et al., 2022). It is very intriguing to speculate that condensate formation will be additionally suppressed in the closed conformation by limiting the accessibility of CFR because only one amino acid separates the identified crucial phenylalanine residues of CFR from the PDZ binding motif in the DVL2 C-terminus. Notably, Wnt ligands induce the open DVL conformation (Harnos et al., 2019), which will increase the CFR accessibility and, according to our model, would allow LCR4 and CD2 to contribute to pre-oligomerization and subsequent condensate formation of DVL2, respectively.

Several studies suggest functional differences between the DVL paralogs (Gentzel et al., 2015; Lee et al., 2008; Paclikova et al., 2021), while the underlying molecular differences remain unclear. Notably, complex formation as revealed by ultracentrifugation was only observed for DVL2 and not for DVL1 or DVL3, revealing a marked difference between the paralogs (Fig. 1C). In addition, CFR did not co-localize with DVL1 or DVL3 in contrast to DVL2, indicating that CFR-mediated DVL2-DVL2 interaction, which most likely drove DVL2 complexation, is absent from DVL1 and DVL3 (Figs. 5B and S5H). Consistently, the crucial CFR aggregon located in LCR4 was not conserved in DVL1 or DVL3 (Fig. S1D). Moreover, replacing the complementary part of DVL1 with the DVL2 CFR promoted complex formation (Fig. 3H, I). Although DVL1 and DVL3 lack the discovered CFR, all three DVL paralogs are able to form condensates (Fig. 2D and S5H) and to activate Wnt signaling to some extent (Lee et al., 2008; Paclikova et al., 2021), which can be potentially explained by the other interaction sites (DIX, DEP, intrinsically disordered region). However, quantitative studies suggest functional differences between the paralogs and that they have to cooperate at a certain molar ratio for optimal Wnt pathway activation, with DVL2 being the most abundant (Lee et al., 2008; Paclikova et al., 2021). In this context, the DVL2 condensates with their underlying stable complexes may function as a kind of superscaffold for integration of DVL1 and DVL3. Our findings may help to understand the functional differences between the paralogs in the future.

## MATERIALS AND METHODS

### Cell culture, transfections and treatments

HEK293T, HeLa and U2OS cells were grown in low glucose DMEM supplemented with 10% fetal calf serum and antibiotics at 37°C in a 10% CO2 atmosphere, and passaged according to ATCC recommendations. T-REx *DVL1/2/3* triple knockout cells (DVL tKO) and T-REx *DVL1/2/3*, *RNF43* and *ZNRF3* penta knockout cells (DVL tKO+) were generated in the Bryja lab and have been previously described (Paclikova et al., 2017; Paclikova et al., 2021). They were grown in high glucose DMEM supplemented with GlutaMAX, 10% fetal calf serum and antibiotics at 37°C in a 10% CO2 atmosphere. Transfections of plasmids and siRNA were performed with polyethylenimine and Oligofectamine (Invitrogen) according to manufacturer’s recommendations, respectively. Wnt3a medium was prepared as originally described (Willert et al., 2003). For the osmotic shock treatment, cell culture medium was diluted 1:1 with sterile water to reduce the osmolarity by 50%. 1,6-Hexandiol (240117) was obtained from Sigma-Aldrich.

### Molecular Biology

The expression vectors for HA-DVL1, HA-DVL2, HA-DVL3 and HA-DVL2 M2 have been described previously (Bernkopf et al., 2015). The expression vectors for HA-ΔDEP, HA-ΔDEP-ΔCD1, HA-ΔDEP-ΔCD2, HA-ΔDEP-ΔLCR1, HA-ΔDEP-ΔLCR2, HA-ΔDEP-ΔLCR3, HA-ΔDEP-ΔLCR4, HA-ΔDEP-ΔCFR, HA-ΔDEP-ΔCFR+CFR^Ax^, HA-1-418, HA-1-418+CD2, HA-1-418+LCR4, HA-1-418+CFR, HA-DVL1-CFR^DVL2^, HA-DIX, HA-DIX+CFR, HA-ΔCFR, Flag-DAX, Flag-CFR+DAX, Flag-1xCFR^Ax^-DAX, Flag-2xCFR^Ax^-DAX, Flag-3xCFR^Ax^-DAX, Flag-CFR were cloned via standard molecular biology methods. HA-1-418+CFR M2, HA-DIX+CFR M2, Flag-CFR+DAX M3, Flag-3xCFR^Ax^-DAX M3, HA-ΔDEP VV-AA, HA-ΔDEP FF-AA, HA-1-418+LCR4 VV-AA, HA-1-418+CD2 FF-AA, Flag-CFR FF-AA, Flag-CFR VV-AA FF-AA, HA-DVL2 VV-AA FF-AA were generated using site-directed mutagenesis. All generated expression vectors were verified by sequencing.

### Antibodies and siRNA

We used the following antibodies in this study: Primary antibodies:: rb α DVL2 [WB: 1:1000], 3216S; rb α DVL2 [WB: 1:1000], 3224S; rb α Axin1 [WB: 1:1000], 2087S CellSignaling / rat α HA [WB: 1:1000], 11867423001 Roche / rat α α-tubulin [WB: 1:1000], MCA77G Serotec / m α Flag [IF: 1:800], F3165; rb α Flag [IF: 1:300], F7425; rb α HA [IF: 1:200], H6908 Sigma-Aldrich. Secondary antibodies: goat α mouse/rabbit-Cy3 [1:300], goat α rabbit-Cy2 [1:200], goat α mouse/rabbit/rat-HRP [1:2000] (Jackson ImmunoResearch). The siRNA targeting human DVL2 (5’-GGAAGAAAUUUCAGAUGAC-3’) was published (Soh and Trejo, 2011).

### Sucrose gradient ultracentrifugation

Cells were lysed about 24 h after transfection, when required, or 48 h after seeding in a Triton X-100-based buffer (150 mM NaCl, 20 mM Tris-HCl pH 7.5, 5 mM EDTA, 1% Triton X-100, Roche protease inhibitor cocktail). A linear sucrose density gradient was prepared in 13×51 mm centrifuge tubes (Beckman Coulter) by overlaying 2 ml of a 50% (v/w) sucrose solution with 2 ml of a 12.5% (v/w) sucrose solution followed by horizontal incubation of the tube for 3 h at RT (Stone, 1974), bevor loading a 200 µl cell lysate sample on top. After centrifugation in a Beckman Coulter Optima MAX Ultracentrifuge (217100 g, 25°C, 18 h), 20 fractions à 200 µl were collected from top to bottom and analyzed by Western blotting, as indicated. The commercially available size markers thyroglobulin (669 kDa, T9145) and albumin (66 kDa, A8531) were obtained from Sigma Aldrich. In case of thyroglobulin and albumin, fractions were analyzed by silver staining of the proteins in polyacrylamide gels.

### Western Blot

Proteins in cell lysates or in fractions of sucrose gradients were denatured, separated by gel electrophoresis in polyacrylamide gels under denaturing conditions (SDS-PAGE), and transferred onto a nitrocellulose membrane (VWR). The proteins were detected using suitable combinations of primary and HRP-conjugated secondary antibodies (see above) via light emission upon HRP-catalyzed oxidation of luminol in a LAS-3000 with Image Reader software (FUJIFILM). Intensities of protein bands were quantified with AIDA 2D densitometry.

### Immunofluorescence

Cells were fixed in a 3% paraformaldehyde solution, permeabilized with 0.5% Triton X-100 and blocked with cell culture medium to reduce unspecific antibody binding, before proteins of interest were stained with suitable combinations of primary and fluorochrome-conjugated secondary antibodies (see above). Analysis and image acquisition was performed at an Axioplan II microscope system (Carl Zeiss) using a Plan-NEOFLUAR 100x/1.30 NA oil objective and a SPOT RT Monochrome camera (Diagnostic Instruments). Cells were categorized in a blinded fashion as “cell with condensates” when exhibiting more than three distinct sphere-like structures, to reduce the number of false positives. The Spot Detector tool of the Icy open source bio-imaging software (Institut Pasteur, version 2.2.1.0) was used to objectively quantify numbers of condensates per cell (Olivo-Marin, 2002).

### Live-cell imaging

For live-cell imaging of the osmotic shock treatment, the culture medium of cells expressing indicated GFP-tagged proteins was replaced with a 50% hypoosmolar phosphate buffered saline solution. Images were acquired at constant exposure times every 15 s over the next 3 min at an Axiovert25 microscope system (Carl Zeiss) using a LD A-Plan 40x/0.50 Ph2 objective and a SPOT Insight QE camera with the SPOT Basic software (Diagnostic Instruments, version 4.0.1). Movies were rendered using Photoshop 19.1.5 (Adobe).

### Luciferase reporter assay

Cells were transfected with a luciferase reporter plasmid either with a β-catenin-dependent promoter (TOP, Tcf optimal) or with a β-catenin-independent control promoter (FOP, far from optimal), a constantly active β-galactosidase expression plasmid and expression plasmids for the proteins of interest, as indicated in the figures. After lysis (25 mM Tris-HCl pH 8, 2 mM EDTA, 5% glycerol, 1% Triton X-100, 20 mM DTT), the luciferase activity was measured via light emission upon luciferin decarboxylation in a Centro LB 960 Microplate Luminometer (Berthold technologies) and the β-galactosidase activity was assessed as release of yellow ortho-nitrophenol upon ortho-nitrophenyl-β-galactoside hydrolysis using a Spectra MAX 190 (Molecular Devices). The luciferase activities were first normalized to the respective β-galactosidase activities to correct for minor variations in transfection efficiency, before the TOP values were normalized to the respective FOP values to correct for unspecific β-catenin-independent changes. TOP/FOP reporter assays were performed in technical duplicates.

### Statistical analysis

Data sets were probed for statistical significance using two-tailed Student’s *t*-tests in a non-paired (Figs. 3D and S2E-G) or paired (all others) fashion, depending on the experimental setup. Statistical significance is indicated by asterisks in the figures (* p<0.05, ** p<0.01, *** p<0.001), when required, and n values of biological replicates are stated in the figure legends for all experiments. We assumed normal distribution of the data based on the nature of the assays and graphical assessment, which, however, was not formally tested owing to the small sample sizes. P-values for the correlation analyses in Fig. 7H were calculated by the test statistic *t*=*R*\*square root((*n*-2)/(1-*R*^2^)), with n-2 degrees of freedom.

## END NOTES

## Acknowledgements

The authors thank Gabriele Daum for excellent technical assistance. This study was funded by grants from the Deutsche Forschungsgemeinschaft to J.B. (BE 1550/12-1) and D.B.B. (BE 7055/2-1), from the Wilhelm Sander-Stiftung to D.B.B. (2018.017.2), and from the Friedrich-Alexander University Erlangen-Nürnberg Interdisciplinary Center for Clinical Research to J.B. and D.B.B (D30). V.B. was supported by the Czech Science Foundation grant no. GA22-25365S.

## Author contribution

S.A., M.B. and D.B.B. performed the experiments. M.S. performed molecular biology. A.S. and D.B.B. designed the experiments and interpreted the data. P.P. and V.B. provided resources and contributed to data interpretation and manuscript writing. J.B. and D.B.B. conceived the study and wrote the manuscript.

## SUPPLEMENTARY FIGURE LEGENDS

**Fig. S1.**
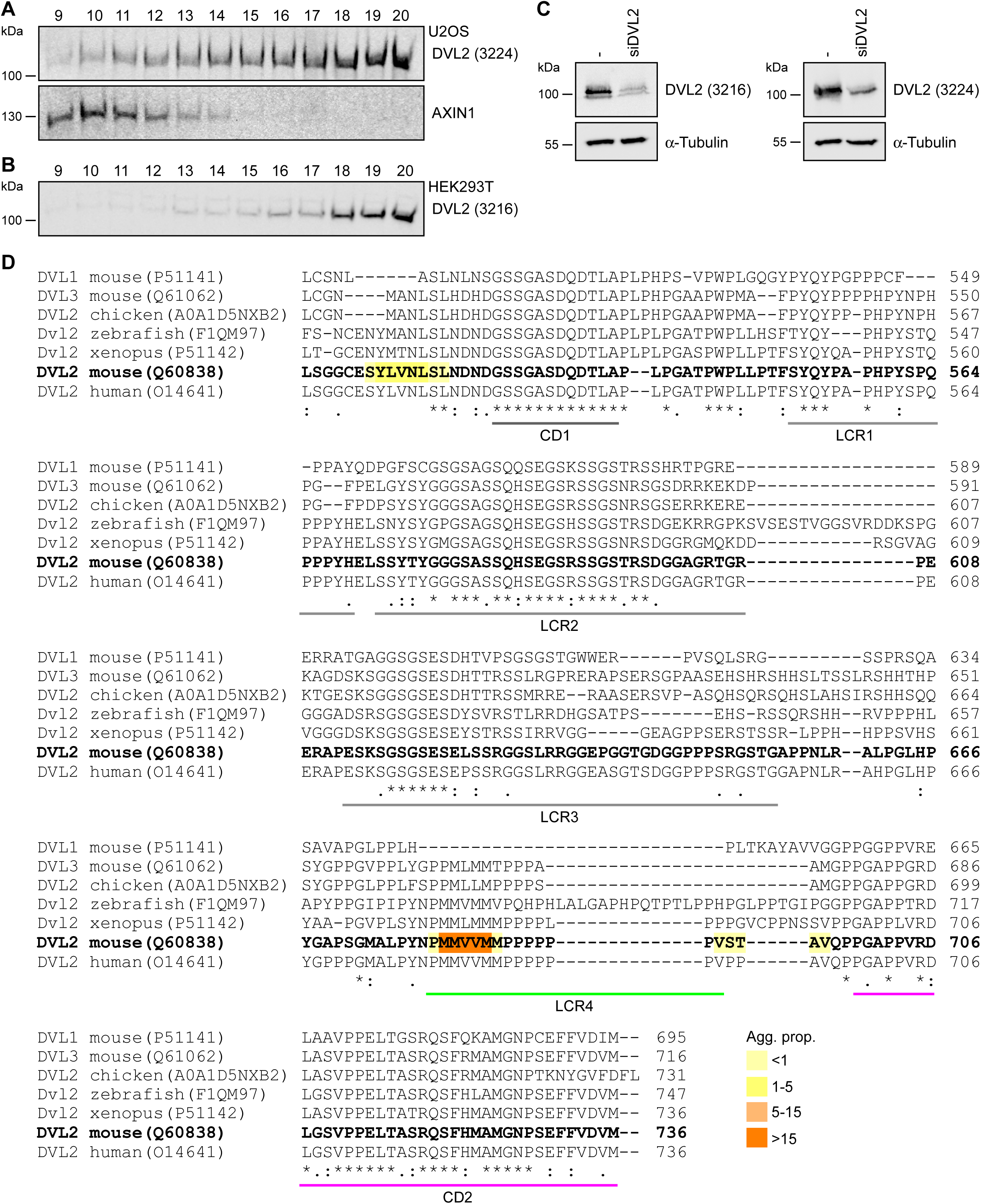
Endogenous DVL2 forms complexes. **A**, **B** Western blotting for indicated endogenous proteins after fractionation of U2OS (**A**) or HEK293T cell lysates (**B**) via sucrose density ultracentrifugation. Fraction numbers are indicated above the blots according to Fig. 1E. **C** Western blotting for indicated endogenous proteins in lysates of HEK293T cells, which were transfected with a control or a DVL2-targeting siRNA for 48 h. **A**-**C** Two different antibodies were used for DVL2 detection (3224 and 3216, CellSignaling). **D** Clustal Omega alignment of indicated dishevelled protein sequences, with specific UniProt identifiers provided in brackets. Identity (*) and conservation between amino acid groups of strongly (:) and weakly (.) similar properties are indicated (Sievers et al., 2011). Marks show conserved domains (CD1 and CD2), low-complexity regions (LCR1, LCR2, LCR3 and LCR4) according to the SEG algorithm (Wootton and Federhen, 1993) and the aggregation propensity of individual residues of mouse DVL2 (yellow-orange heat map) according to the TANGO algorithm (Fernandez-Escamilla et al., 2004). LCR4 and CD2 are highlighted in green and magenta, respectively, as they were identified as crucial regions for condensate induction later on.

**Fig. S2.**
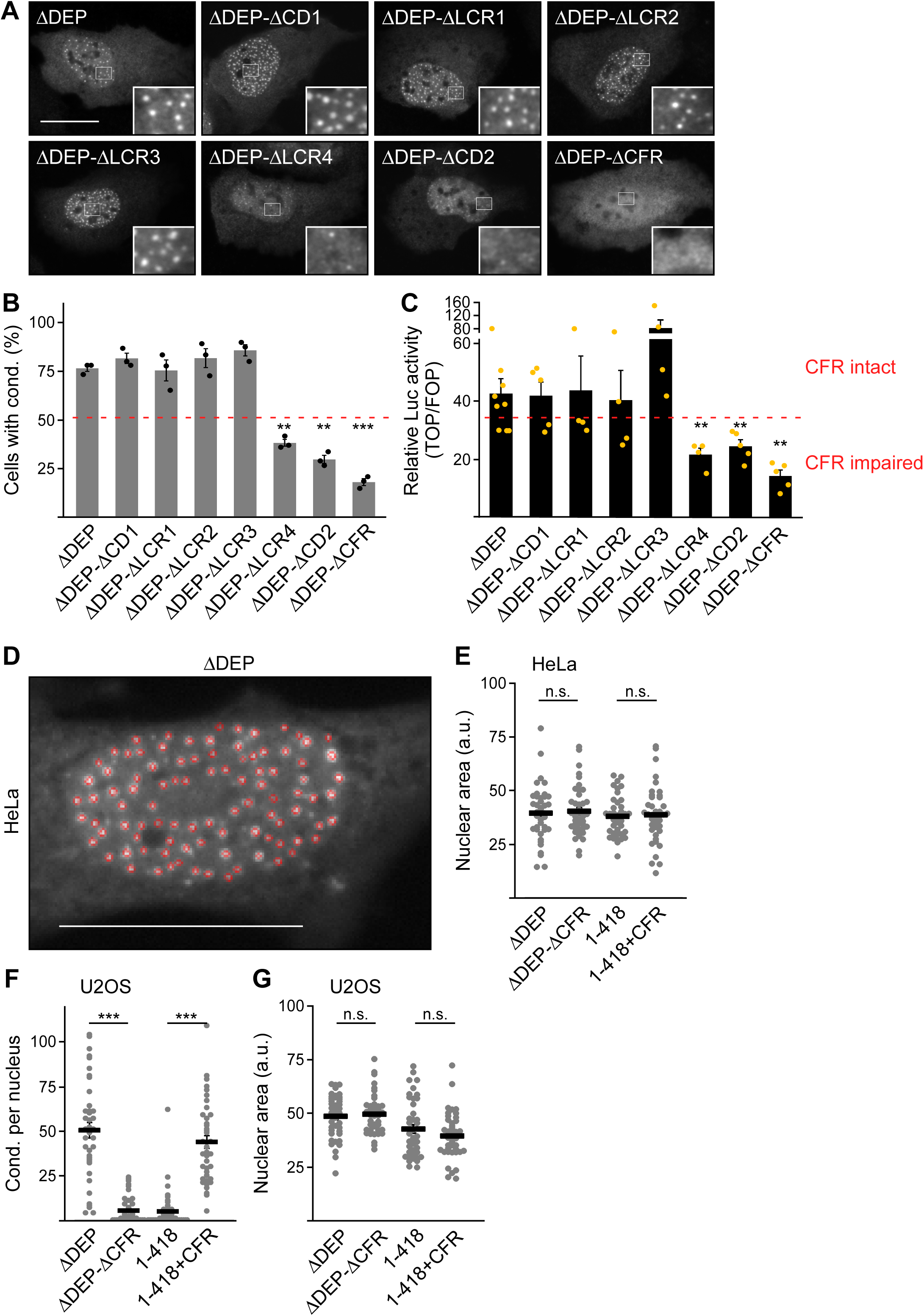
Mapping of DVL2 regions promoting condensates and activity. **A** Immunofluorescence of indicated HA-tagged proteins in transiently transfected U2OS cells. Scale bar: 20 µm. Insets are magnifications of the boxed areas. **B** Percentage of cells with condensates out of 900 transfected cells from three independent experiments as in **A** (n=3). **C** Relative luciferase activity reporting β-catenin-dependent transcription in HEK293T cells expressing the indicated constructs (ΔDEP n=9; ΔCD1, ΔCD2, ΔCFR n=5; ΔLCR1-4 n=4). **B**, **C** We categorized constructs with condensation above and below the dotted red line as CFR intact and CFR impaired, respectively, and this correlated consistently with Wnt pathway activation. **D** Example for automatic detection (red circles) of condensates formed by DVL2 ΔDEP in the nucleus (white signal) using the Icy Spot Detector (Olivo-Marin, 2002), as performed in Fig. 3D. Note the very low number of false negatives (white condensate without red circle) and false positives (red circle without white condensate). Scale bar: 20 µm. **E**, **G** Area of the nuclei analyzed in Fig. 3D (**E**) and in **F** (**G**), showing similar sizes of the investigated nuclei (n=40). **F** Automated quantification of condensate number per nucleus by the Icy Spot Detector in 40 cells from four independent experiments similarly performed as in Fig. 3B but in U2OS cells (n=40). **B**, **C**, **E**-**G** Results are mean +-SEM, ** p<0.01, *** p<0.001 (Student’s *t*-test).

**Fig. S3.**
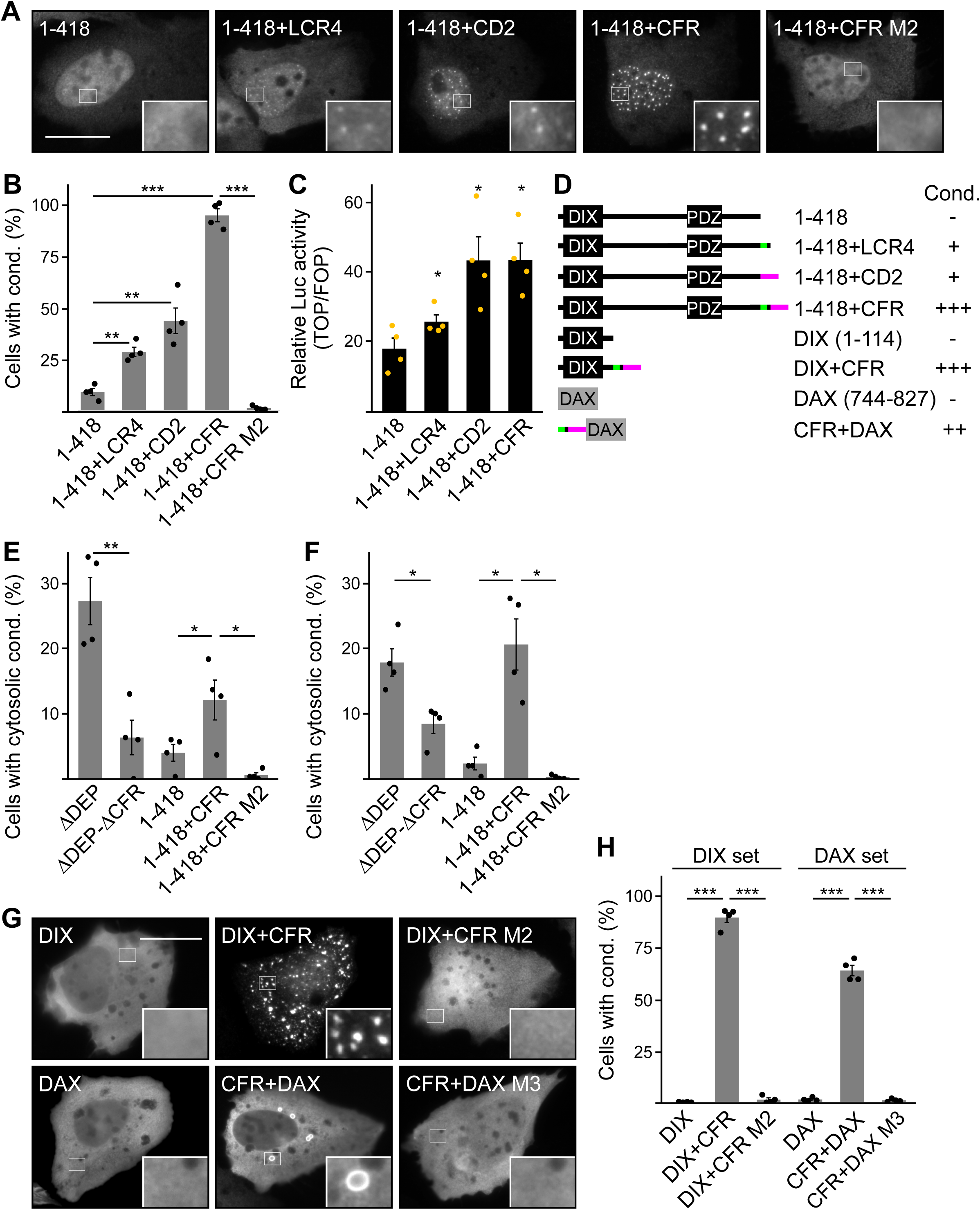
LCR4 and CD2 cooperate to promote DIX domain-dependent condensates. **A**, **G** Immunofluorescence of indicated HA-(**A** and **G** upper row) or Flag-tagged proteins (**G** lower row) in transiently transfected U2OS cells. Scale bar: 20 µm. Insets are magnifications of the boxed areas. **B**, **E**, **F**, **H** Percentage of cells with condensates out of 1200 transfected cells from four independent experiments as in **A** (**B**, **F**), in Fig. 3B (**E**) and in **G** (**H**) (n=4). For **E** (HeLa cells) and **F** (U2OS cells) only cells with cytosolic condensates were quantified. **C** Relative luciferase activity reporting β-catenin-dependent transcription in HEK293T cells expressing the indicated constructs (n=4). **B**, **C**, **E**, **F**, **H** Results are mean +-SEM, * p<0.05, ** p<0.01, *** p<0.001 (Student’s *t*-test). **D** To scale schemes of used constructs with DVL2 and AXIN1 parts depicted in black and grey, respectively. Indicated condensation (Cond.) refers to the findings in **B** and **H**, and the identified crucial regions are highlighted in green (LCR4) and magenta (CD2).

**Fig. S4.**
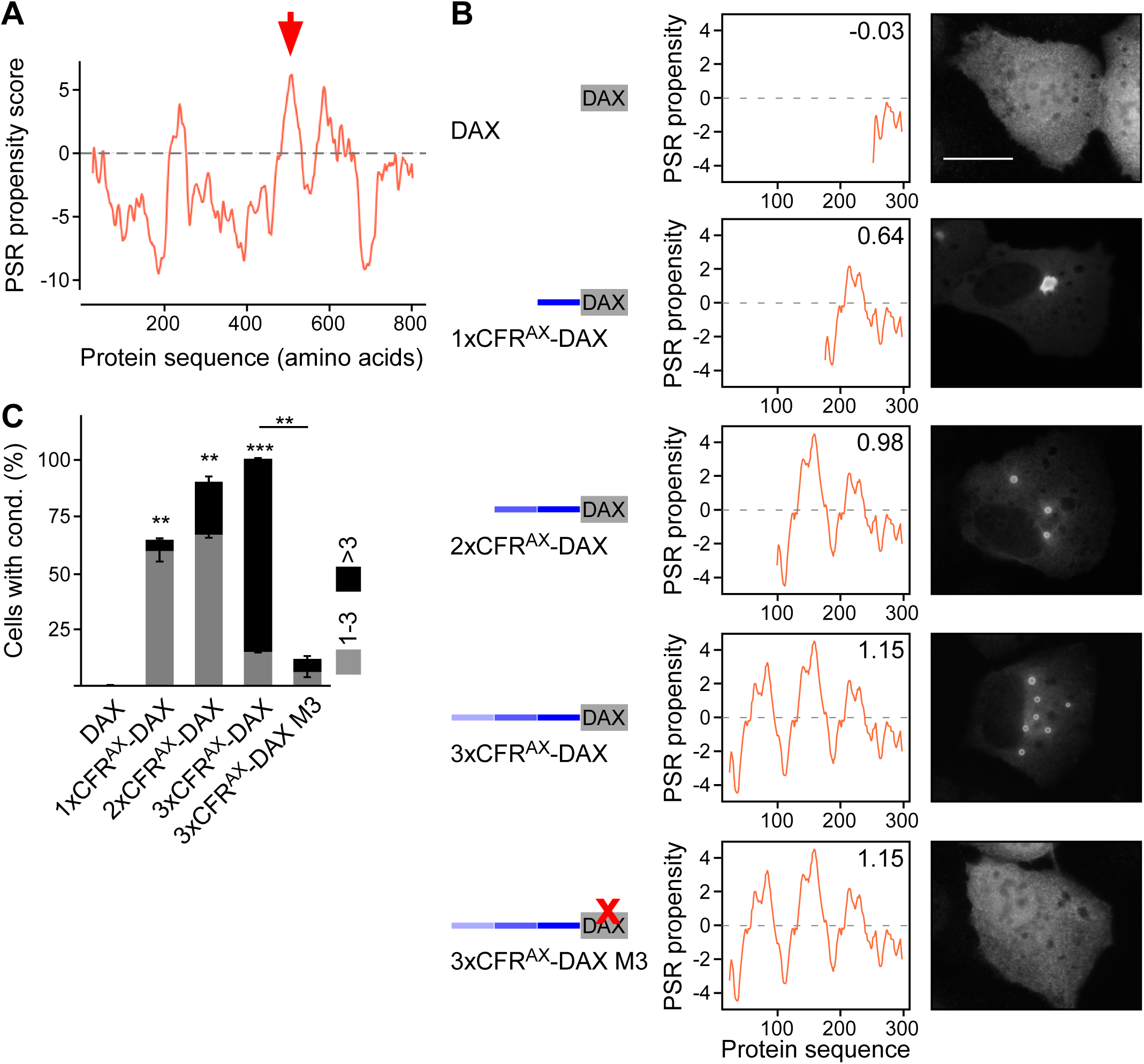
Identifying a phase-separating region in AXIN1. **A** Phase separating (CFR) propensity score (y-axis) of amino acid residues of AXIN1 (x-axis) according to the catGRANULE algorithm, which predicts liquid-liquid phase separation propensity (Bolognesi et al., 2016). Red arrow points to a potential phase separating region in AXIN1 (CFR^AX^). **B** Left column: To scale schemes of artificial fusion proteins between the polymerizing AXIN1 DAX domain (grey) with CFR^AX^ (different shades of blue). CFR^AX^ was fused up to three times to the DAX domain (3xCFR^AX^-DAX). M3 indicates an inhibiting point mutation of DAX-mediated polymerization, which is illustrated by the red X (Fiedler et al., 2011). Middle column: Phase separating (CFR) propensity of the artificial AXIN1 proteins illustrated on the left according to the catGRANULE algorithm. The cumulative CFR scores of the proteins is shown in the upper right corner. Right column: Immunofluorescence of the Flag-tagged proteins indicated on the left in transiently transfected U2OS cells. Scale bar: 20 µm. **C** Percentage of cells with one to three (grey) or more than three condensates (black) out of 900 transfected cells from three independent experiments as in **B** (n=3). Results are mean +-SEM, ** p<0.01, *** p<0.001 (Student’s *t*-test). The increase of condensates upon iterative fusion of CFR^AX^ to the AXIN1 DAX domain validated its activity. Reduced condensation of 3xCFR^AX^-DAX M3 compared to 3xCFR^AX^-DAX demonstrated dependency of condensates on DAX-mediated polymerization, which we used as a specificity control for our artificial constructs, as it is the same for WT AXIN1 (Fiedler et al., 2011).

**Fig. S5.**
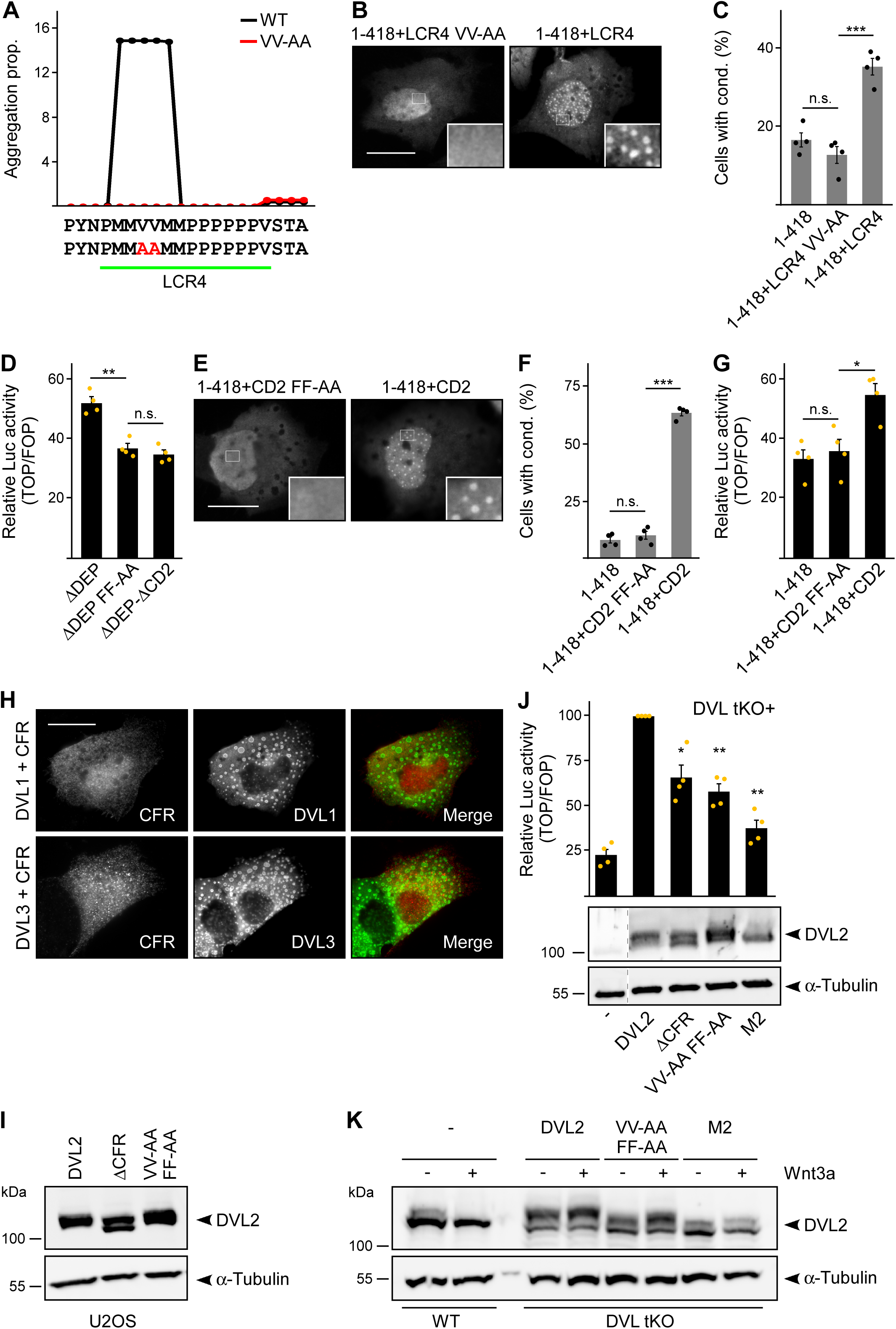
Specifying the residues mediating LCR4 and CD2 function. **A** Aggregation propensity (y-axis) of amino acid residues (x-axis) of WT (black curve) or VV-AA mutated (red curve) DVL2 LCR4 according to the TANGO algorithm, which predicts aggregation nucleating regions (Fernandez-Escamilla et al., 2004). **B**, **E** Immunofluorescence of indicated HA-tagged proteins in transiently transfected U2OS cells. Scale bars: 20 µm. Insets are magnifications of the boxed areas. **C**, **F** Percentage of cells with condensates out of 1200 transfected cells from four independent experiments as in **B** (**C**) and in **E** (**F**) (n=4). **D**, **G**, **J** Relative luciferase activity reporting β-catenin-dependent transcription in HEK293T cells (**D**, **G**) and in T-REx cells with *DVL1/2/3*, *RNF43* and *ZNRF3* knockout (**J** upper panel, DVL tKO+) expressing the indicated constructs (n=4). **C**, **D**, **F**, **G**, **J** Results are mean +-SEM, * p<0.05, ** p<0.01, *** p<0.001 (Student’s *t*-test). **H** Immunofluorescence of indicated proteins in U2OS cells, which were transfected with Flag-CFR together with either HA-DVL1 or HA-DVL3. Scale bar: 20 µm. **I**, **J**, **K** Western blotting showing the expression levels of endogenous DVL2 (-), transiently expressed HA-tagged DVL2 WT (DVL2) and indicated DVL2 mutants to Fig. 7E (**I**), to the upper panel of **J** (**J** lower panel) and to Fig. 7G (**K**) with α-tubulin serving as loading control. One out of three representative experiments is shown. Dotted vertical lines in **J** indicate splicing of the blots.

## SUPPLEMENTARY MOVIE LEGENDS

**Movie M1 Osmotic shock dissolves DVL2 condensates.** Fluorescence of GFP-DVL2 in transiently transfected, alive U2OS cells, which were imaged every 15 s over three min of osmotic shock treatment.

**Movie M2 Osmotic shock dissolves CFR-induced condensates.** Fluorescence of GFP-1-418+CFR in transiently transfected, alive U2OS cells, which were imaged every 15 s over three min of osmotic shock treatment.

